# Diversity of the bacterial and viral communities in the tropical horse tick, *Dermacentor nitens* in Colombia

**DOI:** 10.1101/2023.05.04.539352

**Authors:** Andres F. Holguin-Rocha, Arley Calle-Tobon, Gissella M. Vásquez, Helvio Astete, Michael L. Fisher, Alberto Tobon-Castano, Gabriel Velez-Tobon, L. Paulina Maldonado-Ruiz, Kristopher Silver, Yoonseong Park, Berlin Londono-Renteria

## Abstract

Ticks are obligatory hematophagous ectoparasites that transmit pathogens among various vertebrates, including humans. The composition of the microbial and viral communities in addition to the pathogenic microorganisms is highly diverse in ticks, but the factors driving the diversity are not well understood. The tropical horse tick, *Dermacentor nitens*, is distributed throughout the Americas and it is recognized as a natural vector of *Babesia caballi* and *Theileria equi*, the causal agents of equine piroplasmosis. We characterized the bacterial and viral communities associated with partially-fed *D. nitens* females collected by a passive survey on horses from field sites representing three distinct geographical areas in Colombia (Bolivar, Antioquia, and Cordoba). RNA-seq and sequencing of the V3 and V4 hypervariable regions of the 16S rRNA gene were performed using the Illumina-Miseq platform. A total of 356 operational taxonomic units (OTUs) were identified, in which the presumed endosymbiotic Francisellaceae/*Francisella* spp. was predominantly found. Nine contigs corresponding to six different viruses were identified in three viral families: Chuviridae, Rhabdoviridae, and Flaviviridae. Differences in the relative abundance of the microbial composition among the geographical regions were found to be independent of the presence of *Francisella*-Like Endosymbiont (FLE). The most prevalent bacteria found on each region were *Corynebacterium* in Bolivar, *Staphylococcus* in Antioquia, and *Pseudomonas* in Cordoba. *Rickettsia*-like endosymbionts, mainly recognized as the etiological agent of rickettsioses in Colombia were detected in the Cordoba samples. Metatranscriptomics revealed 13 contigs containing FLE genes, suggesting a trend of regional differences. These findings suggest regional distinctions among the ticks and their bacterial compositions.

## 1 Introduction

Ticks are important vectors of pathogens that cause livestock and human diseases, such as ehrlichiosis, borreliosis, Lyme disease, human and cattle babesiosis, and theileriosis. Tick-borne encephalitis virus, Powassan virus, and Crimean-Congo hemorrhagic fever virus are one of the most prevalent tick-borne viral infections.(1,2). The risks of emerging and re-emerging tick-borne diseases remain a continuing threat since prevention and management are hampered by suboptimal diagnostics, lack of treatment options for emerging pathogens, and scarcity of vaccines (3,4). Habitat changes of the ticks by human activities and globalization have been described as direct factors driving migration and colonization of hosts, vectors, and pathogens (5). In addition, global climate change caused by human activities has increased the incidence and diversity of circulating pathogens in new habitats (6).

Ticks harbor diverse microorganisms, including symbionts, in addition to pathogenic organisms, which may have direct positive/negative effects on the tick or other members of the microbial communities (1,7,8). Interactions among the microorganisms in the bacterial communities in the ticks are considered an important factor in the transmission of human/animal pathogenic organisms. (9,10). Among non-pathogenic communities, common bacterial endosymbionts found in ticks are mainly related to *Rickettsia*, *Coxiella*, and *Francisella* genera (1,11,12). These microorganisms act as primary endosymbionts providing essential nutrients involved in survival, development, and tick-fitness, such as biosynthesis of B vitamins and cofactors like riboflavin, folic acid, and biotin (13). Tick-endosymbionts are generally tissue-specific with microbial guilds well established in salivary glands, gut, ovaries, among other tissues (14). Some of these microorganisms, including pathogenic and non-pathogenic bacteria, can be transovarially transmitted to tick offspring (15). Given the importance of ticks as vectors of many important pathogens, understanding ticks and their symbiont compositions in different ecological systems has arisen as an important area of study (2).

The tick microbiome includes communities of viruses, bacteria, protozoa, and fungi (8,14). Recent experimental approaches to characterize the bacterial diversity in various species of ticks used next-generation sequencing (NGS) of the 16S rRNA gene sequence amplicons (16–18). Those studies revealed tick bacterial communities, including mammalian pathogens, that are dependent on the tick species, type of host, and geographic location (4,11,19). Characterizing the microbial tick populations may give us a better understanding of the different potential roles in intra- and interspecific microbial interactions and their involvement in vector competence (4,7,20).

Viruses are present in all domains of life, particularly rich in the phylum Arthropoda, which includes ticks (21). Metatranscriptomics is a widely used tool to investigate RNA viruses in ticks. Despite considerable insights into bacterial diversity, our understanding of tick-associated viruses is still limited, and largely unexplored compared with bacterial diversity (22). Virome studies of ticks collected in Asia, Europe, and North America have revealed the emergence of novel pathogenic tick-borne viruses as well as the dearth of data on tick viromes which suggest a need for viral surveillance and discovery in this group of arthropods (23–25). Progress in sequencing technology and metagenomics data have provided an approximation to the viral community composition present in a few tick species (22,24,26–30). In addition, more information from different species may be an efficient strategy to mitigate potential threats of tick-borne disease to public health (2,3,25,30).

The tropical horse tick, *Dermacentor nitens*, is distributed throughout the Americas and it is recognized as a natural vector of *Babesia caballi* and *Theileria equi*, the causal agents of equine piroplasmosis (31,32). *Dermacentor nitens* is a one-host tick, with three to four generations per year (33). Severe infestation in vertebrate animals can cause severe lesions, especially in the ears, and predispose the host to secondary bacterial infections (34). Although equines are the primary host, natural infestations have been reported in other domestic, and companion animals, as well as wild animals (35–37). *Dermacentor nitens* is considered a sporadic ectoparasite of humans, where tick infestations are probably a consequence of humans entering infested livestock environments, resulting in a transference of ticks from infested animals to persons (38). Accidental infestations by *D. nitens* in humans related to agricultural activities may represent a potential danger to human health, although the vectorial capacity of *D. nitens* for pathogens related to public health remains unknown. Occurrence of human pathogenic agents in this tick species have been previously reported (39,40).

To gain an in-depth understanding of the microbial communities of *D. nitens*, we used 16S rRNA gene sequences combined with metatranscriptomic analysis to identify the main bacterial and viral communities present in the ticks collected in different geographical populations. These results provide large numbers of sequences annotated as tick viruses and operons of *Francisella*-like endosymbionts (FLE) and revealed a trend of differences among the three geographical populations.

## 2 Materials and methods

### 2.1 Sample collection and nucleic acid extraction

Tick collection was carried out by passive survey at “La Rinconada” slaughterhouse (06°11’26.0”N; 75°22’43.4”W) in the municipality of Rionegro, Antioquia, Colombia in July, and September 2019. A total of 45 blood-fed *D. nitens* adults were obtained from three horses native to each region, Bolivar, Antioquia, and Cordoba (Supplementary Figure 1). The three departments are located in the northwest of Colombia and share borders with the department of Antioquia. Live ticks were transported to the Universidad de Antioquia facilities, where taxonomical identification was made following morphological keys (41), and specimens subsequently stored at −20 or −80°C until shipment to Kansas State University facilities. Blood-fed female *D. nitens* collected from horses were pooled and processed based on host (individual animal) and region (Bolivar, Antioquia, and Cordoba). From a total of three horses per region and one pool of five ticks per horse were chosen by using the random selection method, thus sampling a total of 45 ticks (nine pools). Genomic DNA and RNA were extracted independently following manufacturer instructions using Zymo™ DNA and RNA extraction kits (Irvine, California, US) from the pools previously separated from the tick-exoskeleton.

### 2.2 NGS library preparations and data processing

Genomic DNA of the pools of ticks was sent to the Genome Sequencing Core at the University of Kansas. Amplicon libraries were prepared by Illumina Miseq targeting the V3-V4 region with the primers 16S-F (5’-TCGTCGGCAGCGTCAGATGTGTATAAGAGACAGCCTACGGGNGGCWGCAG-3’) and 16S-R (5’-GTCTCGTGGGCTCGGAGATGTGTATAAGAGACAGGACTACHVGGGTATCTAATCC-3’) of the 16S rRNA, with an expected length of ∼465 base-pair (bp) for the DNA analysis (16).

16S rRNA sequences were analyzed with Mothur v.1.45, according to the MiSeq Standard Operating Procedure (42). Operational Taxonomic Units (OTUs) with 97% of identity were clustered and classified using the database SILVA v.138. Raw reads were filtered to a maximum length of 465 base-pair without ambiguous bases (43). Another filtering step was done in Excel to remove low-count OTUs with a prevalence in samples of less than 0.005% (44). Bacterial relative abundance was analyzed in R studio (vegan and ggpubr packages), and GraphPad Prism 9.2.0 software (45–47). We also compared the differences in the proportion of the bacterial composition of the regions through a Non-Metric Multidimensional Scaling (NMDS) ordination plot. It is important to note that there is the potential for low-frequency background noises in this dataset due to the absence of blank extraction control during the nucleic acid extraction and bioinformatics workflows (44).

RNA-seq library preparation was done with the NEB Next Stranded RNA library kit without PolyA selection of the mRNA, the nine pooled RNAs were sent to the Genome Sequencing Core at the University of Kansas. For the metatranscriptomics analysis, the RNA-seq reads were processed for removal of Illumina adaptor sequences, trimmed, and quality-based filtered using Fastp software v.0.20.0 (48). The high-quality reads (Phred-score >30) were removed by mapping onto the reference genome of *D. silvarum* (assembly ASM1333974v1) and *Equus caballus* (assembly EquCab3.0) using STAR v.2.7 (49). The unmapped reads (Supplementary Table 1) were used to perform the assembly and annotation of the transcriptome by using Trinity and Blast2GO suite in OmicsBox v.2.0.36 software (50–52). Contigs annotated in Blast2GO were reexamined manually by BLASTn and BLASTx (https://blast.ncbi.nlm.nih.gov/Blast.cgi) to confirm the results and eliminate potential false positives. Empirical Bayes estimation and Fisher’s exact tests (α = 0.05) by pairwise comparison based on the negative binomial distribution analysis were done with edgeR by using the Galaxy platform to test statistically significant differences in abundance between the bacterial and viral sequences annotated with the geographic location for the blood-fed *D. nitens*.

### 2.3 Phylogenetic analyses of viral and *Francisella* spp. contigs

Phylogenetic analyses by comparison of Bayesian inference, Maximum-Likelihood, Minimum-Evolution, and Neighbor-Joining methods were performed as an initial assessment with the bacterial protein sequences and the OTUs detected in this study compared to the reference sequences pulled out from the NCBI GenBank database by doing homology-based search using Blast search. Bacterial protein sequences, partial 16s rRNA nucleotide sequences of FLE, and viral protein sequences were retrieved from the GenBank database as indicated with the GenBank accession numbers in Figures 2 to 4. Sequences were aligned by using Muscle in MEGA-X software (53). Bayesian inference analysis was done using BEAST v1.10.4 software (54). Phylogenetic trees for the analysis of the 16s rRNA nucleotide sequences were constructed based on the Neighbor-Joining method with a pairwise deletion. The tree for the V3-V4 regions sequenced in this study were constructed with 500 bootstrap replicates (55–57) unless otherwise specified. For metatranscriptomic analyses of the FLE and viral proteins sequences, the cladograms were constructed using annotated and concatenated genes for each contig by using the Maximum Likelihood method with Tamura-Nei model and 500 bootstrap replicates (58).

### 2.4 Ethical approval

This study was approved by the Bioethics Committee of the Universidad de Antioquia (Approval record No. 15-32-436 of June 2015). It was also granted an environmental license issued by the Colombian government through the National Environmental Licensing Authority (Autoridad Nacional de Licencias Ambientales-ANLA, Resolution ANLA 00908 of May 27, 2017).

## 3 Results

### 3.1 Bacterial diversity investigated using V3-V4 regions of the 16S rRNA sequences

A total of 372,493 sequences after filtering 392,819 raw reads were assembled into 6,686 contigs and assigned to 356 OTUs with a threshold of 97% of sequence identity (Table 1). Notably, the sequences consisted of three main OTUs, all identified as FLE (>80%) in all nine samples (Figure 1A). Among the remaining <20% OTUs, the most prevalent bacteria in different regions were *Corynebacterium* in Bolivar, *Staphylococcus* in Antioquia, and *Pseudomonas* in Cordoba (Figure 1B). We also compared the differences in bacterial compositions of the regions through Non-Metric Multidimensional Scaling (NMDS) in the data sets before and after excluding FLE (Figures 1C and 1D). Our NMDS plots suggest that regional bacterial composition is unique and independent of the presence of FLE and can be useful to differentiate the bacterial composition from different geographical regions (Figure 1).

**Figure 1.**
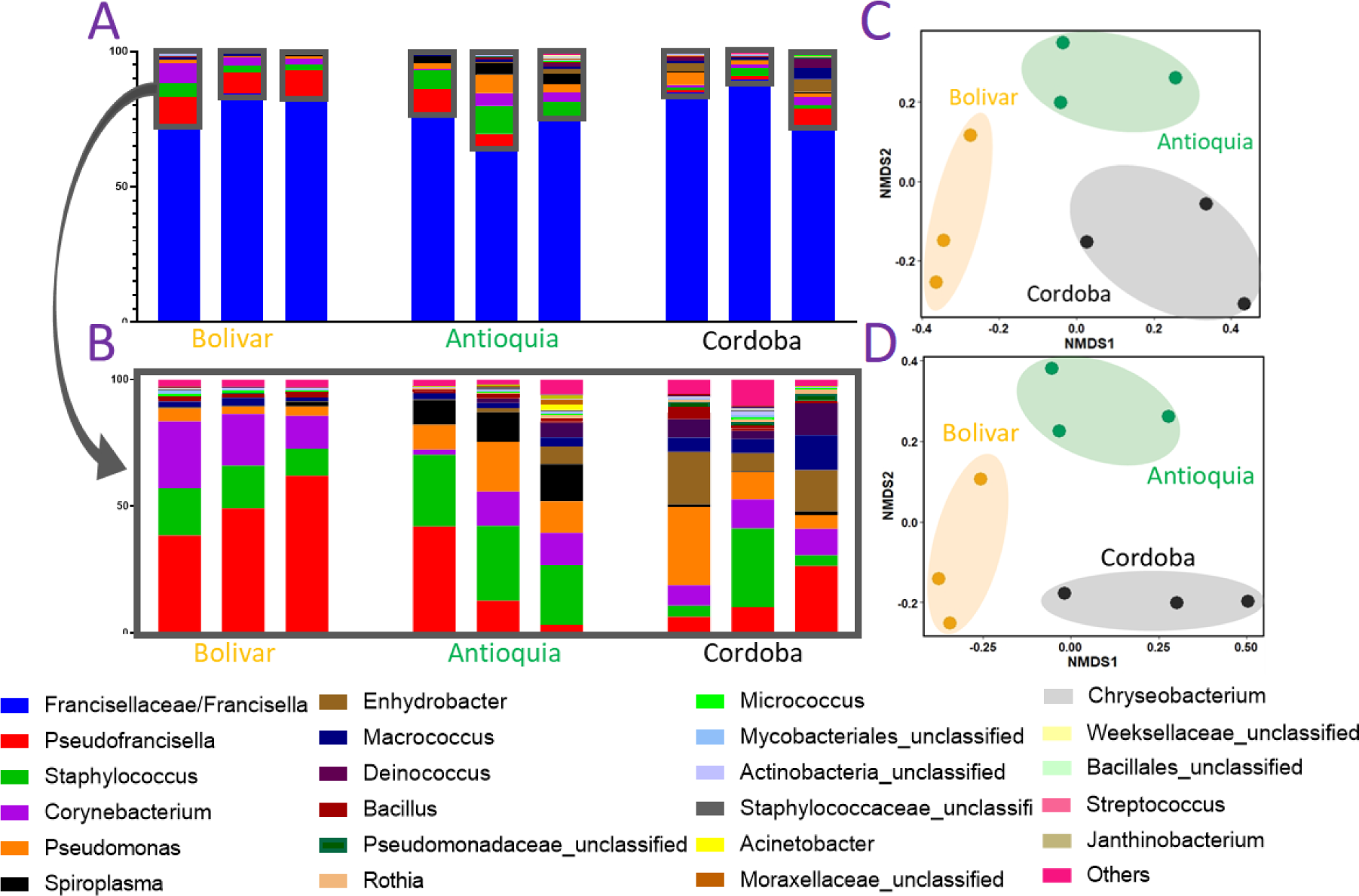
Bacterial diversity shown by the genera in 16S rDNA sequences from *Dermacentor nitens* samples collected from three different regions of Colombia. **(A)** Relative abundance is shown by bacterial genera. **(B)** The relative abundance after excluding the sequences of endosymbionts Francisellaceae/*Francisella* spp. **(C)** Non-metric multidimensional scaling plot (NMDS) plot showing the differences among tick samples from different regions. **(D)** NMDS plot showing the differences among tick samples after excluding the endosymbionts.

**Table 1.** Nine sequencing libraries for the pools for *D. nitens,* targeting V3-V4 regions of the 16 rRNA gene.

| Library (Paired Reads) | Region | Raw reads | Mapped Reads | Contigs |
| --- | --- | --- | --- | --- |
| DNA_Pool_1 | Bolivar | 48852 | 46109 | 706 |
| DNA_Pool_2 | Bolivar | 41430 | 39512 | 508 |
| DNA_Pool_3 | Bolivar | 37846 | 36438 | 503 |
| DNA_Pool_4 | Antioquia | 45141 | 42948 | 842 |
| DNA_Pool_5 | Antioquia | 39380 | 37847 | 1044 |
| DNA_Pool_6 | Antioquia | 43778 | 41116 | 886 |
| DNA_Pool_7 | Cordoba | 47878 | 45604 | 665 |
| DNA_Pool_8 | Cordoba | 41244 | 38268 | 583 |
| DNA_Pool_9 | Cordoba | 47270 | 44651 | 949 |
| Total |  | 392819 | 372493 | 6686 |

The FLEs categorized by a 97% identity threshold were three different OTUs (OTU001, 002, and 010 in Figure 2A and Supplementary Table 2). These sequences are significantly different from each other with 20 nucleotides (nt) mismatches between OTU001 and OTU002, 21 nt mismatches between OTU002 and OTU010, and 8 nt mismatches between OTU001 and OTU010. High frequencies of the reads for each FLE OTUs, which are in independent libraries, suggest that the three different FLE OTUs are not sequencing artifacts. The cladogram of the FLE sequences showed these three OTU clustered in a branch with the bootstrapping value of 100 (Figure 2A). A single OTU, OTU184, was categorized into *Rickettsia*-like endosymbiont (RLE) in one pool of the Cordoba region. Phylogenetic analysis supports the position of this sequence in the tree clustered with RLE of *Amblyomma latepunctatum* and a clear separation from the pathogenic *Rickettsia* although the bootstrapping value was 68 (Figure 2B).

**Figure 2.**
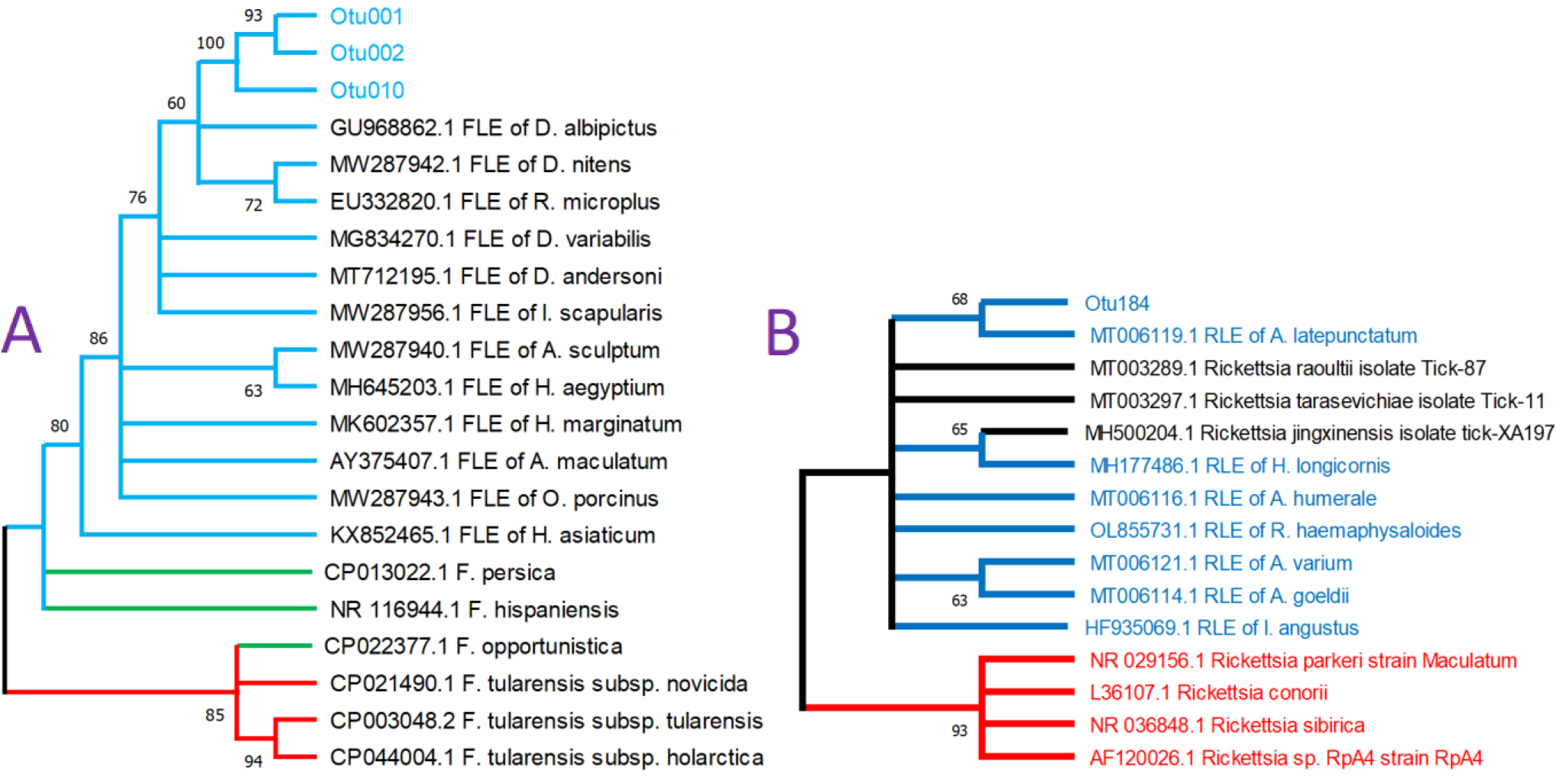
Phylogenetic analyses for the *Francisella-*Like endosymbionts (FLE, **A**) and *Rickettsia*-like endosymbionts (RLE, **B**) identified in this study for *Dermacentor nitens* samples. **(A)** Neighbor-joining cladogram rooted to *Francisella tularensis* strains representing the phylogenetic relationship of 16S rDNA sequences OTUs classified as *Francisella* spp. in *D. nitens*. The tree was built using the pairwise deletion method. The blue branches represent the FLE clade, the green branches represent opportunistic pathogenic *Francisella* species, and the red branches represent the pathogenic *Francisella tularensis* strains as an outgroup. **(B)** Neighbor-joining cladogram rooted to pathogenic *Rickettsia* strains to represent the phylogenetic relationship of rickettsial 16S rDNA sequences with the OTU184 classified as *Rickettsia* spp. in the *D. nitens* sample. The red branches represent pathogenic *Rickettsia* spp., blue branches represent the sequences of RLE, and dark branches represent candidate-human pathogenic *Rickettsia*. The OTUs were determined by a 97% identity threshold. Bootstrapping percentages in 500 replications are shown on the nodes with a 60% cut-off. The GenBank accession numbers for each sequence are shown at the beginning of names of taxa.

### 3.2 Metatranscriptome containing viral and *Francisella* spp. RNA

A total of 152.2 million raw reads were obtained from the nine pools representing the three different regions. After quality trimming and filtering out against *E. caballus* and *D. silvarum* sequences, 92.18 million reads were used for downstream analysis (Supplementary Table 1). *De novo* assembly was conducted using the TRINITY pipeline built in OmicsBox software. After cleaning and filtering, 16.8 million reads were assembled into 81 contigs. Homology-based taxonomic assignment and gene function for each contig was made in Blast2Go and using manual BLAST searches.

Thirteen contigs were categorized as FLE, containing presumed independent operons with an average length of 4,794 bp. Table 2 represents the length and coverage information, the sequence name, the gene encoded, and the putative gene size for each contig (Supplementary Figure 2). The highest coverage of the FLE contigs was Contig_ORF_FLE_of_D. nitens_13, which partially encodes the Mechanosensitive ion channel protein MscS with a length of 596 and 1,892.14 TPM (transcripts per million reads) (Supplementary Figure 3 and Supplementary Table 3). FLE putative operon sequences were submitted to GenBank with the accession numbers contained in the BioProject PRJNA953638.

**Table 2.**
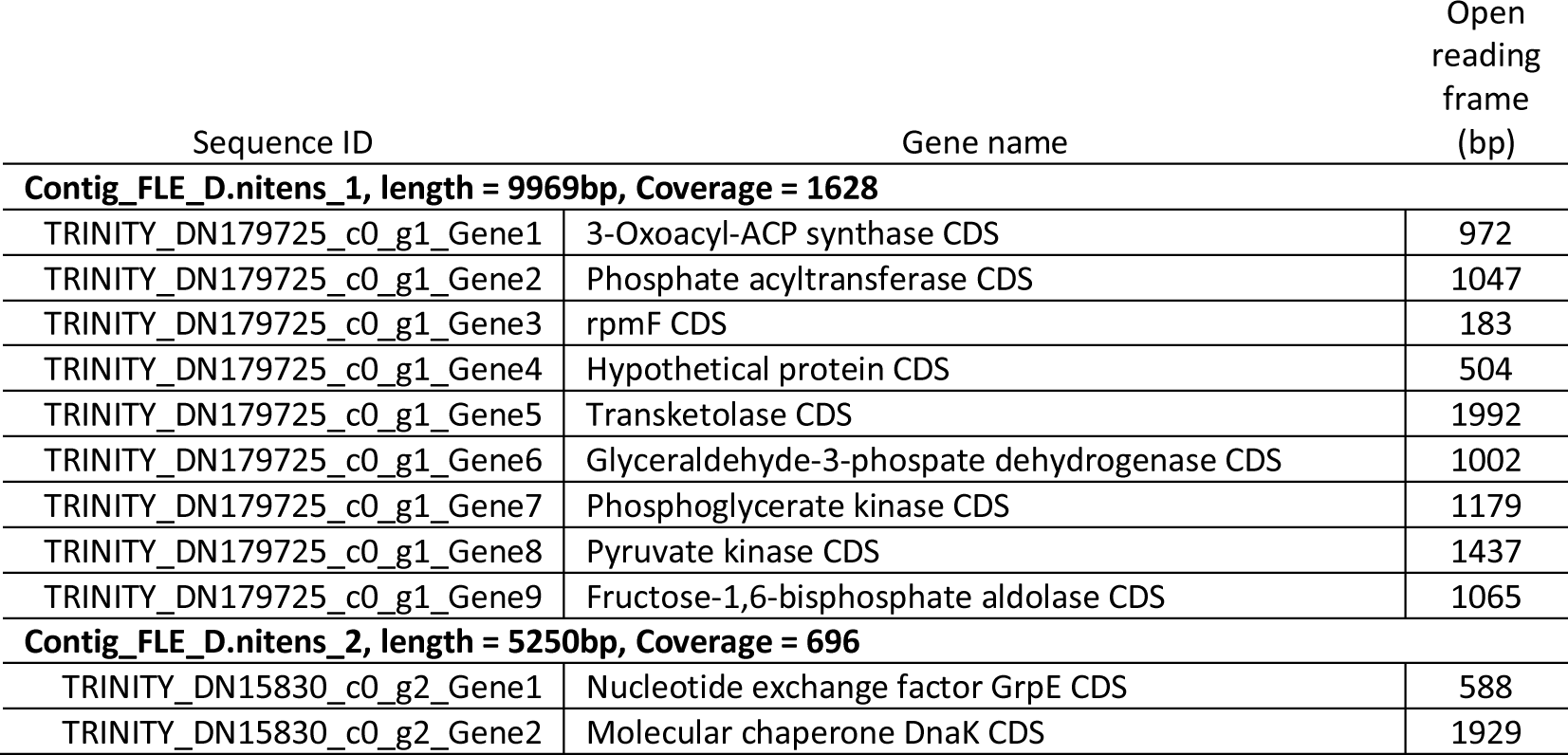
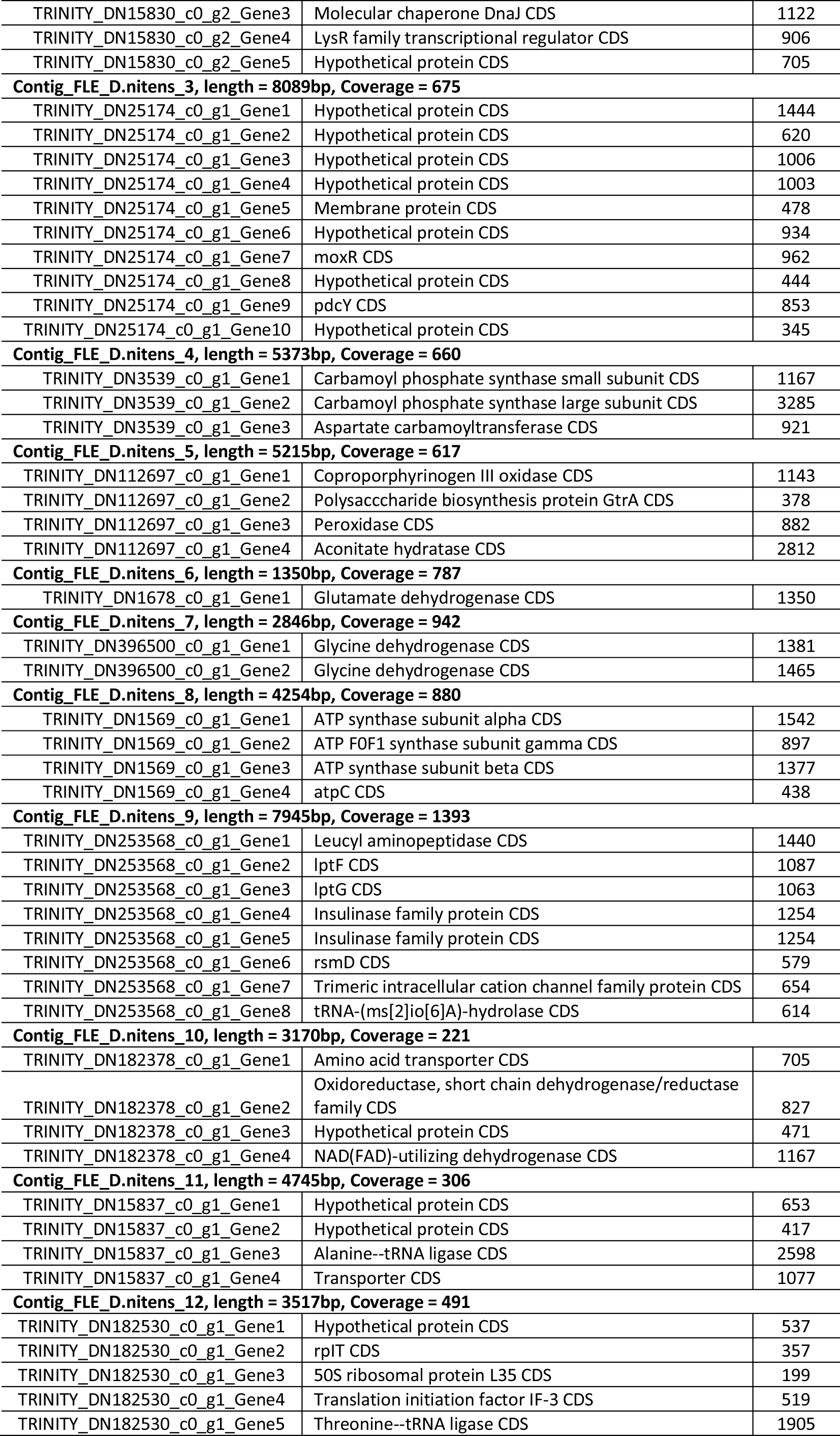
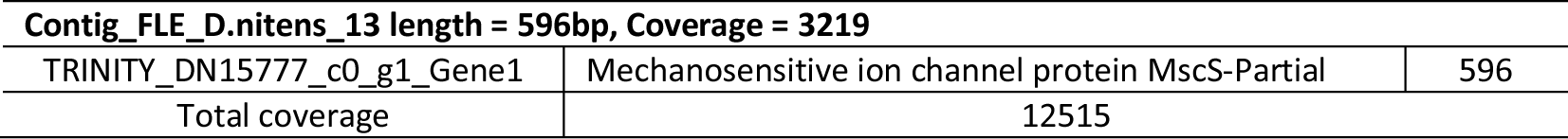
Annotations of bacterial contigs captured in the metatranscriptome of *Dermacentor nitens*.

Six different putative viruses covered by nine viral contigs with an average length of 1,749 bp were identified in BLAST searches for the non-redundant protein database of NCBI and the Viral Genomes database. The sequences were manually inspected and annotated for the coding regions. Table 3 shows the viral contigs with the length and coverage information. The highest coverage for the viral contigs was the D. nitens_Colombia_Flaviviridae_Polyprotein_6 contig with a total of 2,346.25 TPM with the coverage predominantly higher in the region of Cordoba (Supplementary Figure 4 and Supplementary Table 4). The *D. nitens* virus contig sequences were submitted to GenBank with the accession numbers contained in the BioProject PRJNA953638.

**Table 3.**
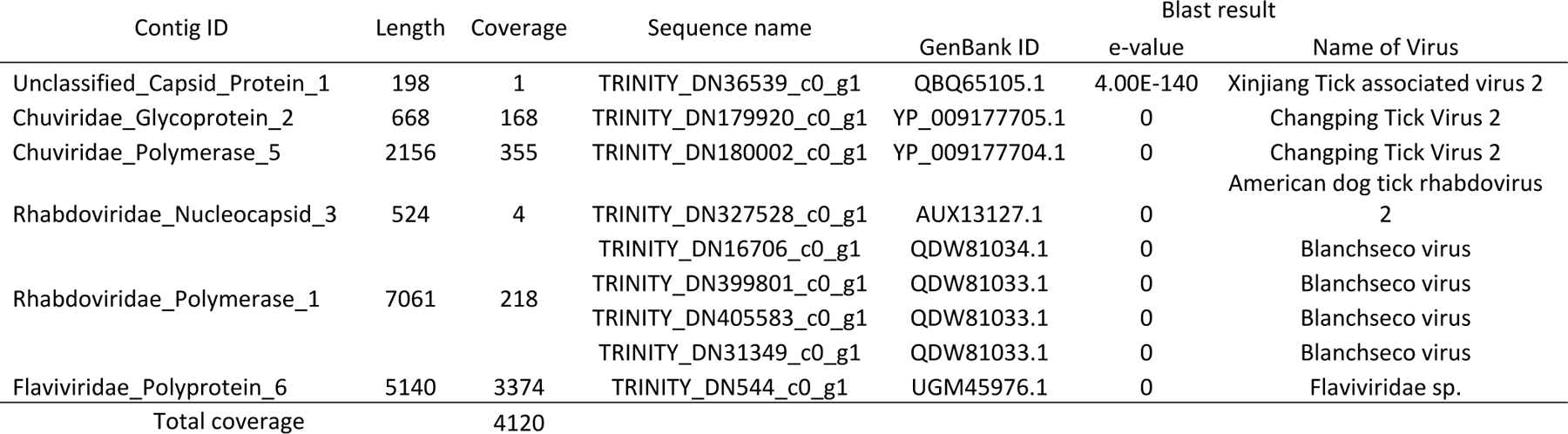
Viral contigs captured in the metatranscriptome of *D. nitens*, shown for the lengths, coverages, and Blast results.

### 3.3 Phylogenetic analyses of viral and *Francisella* spp contigs

Thirteen FLE groups and nine viral contigs identified by metatranscriptomics were further analyzed for their phylogenetic positions. All 13 FLE contigs clustered with other FLE identified in tick species when rooted in the pathogenic and opportunistic *Francisella* groups. The sequences had a 100% bootstrapping value for the tick endosymbiont clade represented by *Amblyomma maculatum* and *Ornithodoros moubata* (59) Figure 3 showing the phylogeny of concatenated sequences of 13 contigs. The overall similarity was 90% with the FLE of the Ixodidae family represented by *A. maculatum*. The green branched clade, containing *F. persica*, *F. opportunistica*, and *F. hispaniensis* represents the opportunistic pathogens that have been linked as potential causative agents of illness episodes in humans (12,59,60). The red branched cluster, shown as the outgroup, are the pathogenic strains of *Francisella tularensis sl*. To show the relationship of the contigs identified with the FLE clade, the sequence named Contig_ORF_FLE_of_D.nitens_1 was used as a representative sequence for the phylogenetic analysis, mainly because all 13 contigs grouped with the tick endosymbiont clade. The total coverage found for the 13 contigs classified as FLE was 12,515, with contigs 13 and 1 being the most predominant among all pools of samples (Supplementary Table 3).

**Figure 3.**
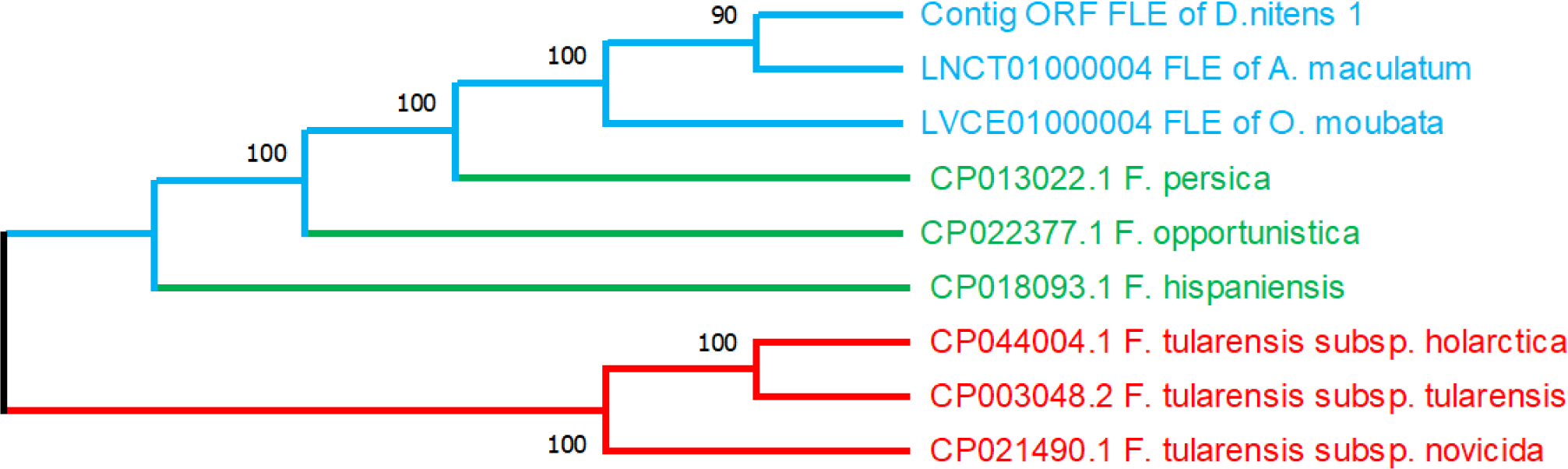
Phylogenetic relationship of the *Francisella*-Like Endosymbiont in the *D. nitens* samples in this study. The sequence is the translated sequence for the concatenated open reading frames. The selected contig contains nine genes (Table 2) annotated with a total length for the concatenated contig of 3323 amino acids (9969 bp). and 1892 transcript per million (TPM) in the pooled metatranscriptome. The tree is for maximum likelihood cladogram built using the complete deletion method. Bootstrapping percentage values are based on 500 replications and are shown at the nodes. The outgroup is for the sequences of pathogenic *F. tularensis* strains. The blue lines correspond to tick FLE, the green lines correspond to opportunistic pathogens, and the red lines correspond to pathogenic strains of *F. tularensis*. The GenBank accession numbers are shown at the beginning of each label.

Phylogenetic analysis of nine viral contigs found three different families for all different viral species. The genes were capsid protein, glycoprotein, nucleocapsid, polyprotein, and RNA-dependent RNA polymerase (RdRp) (Table 3). Most of the putative viruses were found by identifying genes encoding RdRp with five annotated sequences and classified into two viral families, Chuviridae and Rhabdoviridae. Two different contigs, D. nitens_Colombia_Chuviridae_Glycoprotein_2, and D. nitens_Colombia_Chuviridae_RdRp_5 were grouped into the same family Chuviridae. Based on the sequence similarities and the tree pattern (Figures 4A and 4B), these contigs are likely presenting two different viruses although the name of the closely related virus is the same as Changping Tick Virus 2, a virus that has been reported in China and Turkey infecting *Dermacentor* spp. and *Hyalomma asiaticum* ticks (23,24). These two viruses were found to be more abundant in the region of Antioquia (Supplementary Table 4). The Family Rhabdoviridae is represented by five sequences clustered into two putative viruses (Figures 4C and 4D). Four of them targeting RdRp were grouped in a clade with Blanchseco virus. The remaining sequence was found encoding a nucleocapsid protein and clustered with the American dog tick Rhabdovirus-2. The contig D. nitens_Colombia_Unclassified_Capsid_Protein_1 showed a close relationship with the capsid protein of Xinjiang tick-associated virus-2, a virus sequence that was presumably reported for the first time in the province of Xinjiang in China. This virus remains as unclassified for the family, and it is grouped with other tick viruses found in *Ixodes scapularis* and *D. variabilis* (Figure 4E). The family Flaviviridae was found to be represented by one contig named D. nitens_Colombia_Flaviviridae_Polyprotein_6 (Figure 4F). This name was assigned due to the high similarity found with a portion of a Flaviviridae polyprotein from *Haemaphysalis longicornis* and *Rhipicephalus microplus* infesting goats (30).

**Figure 4.**
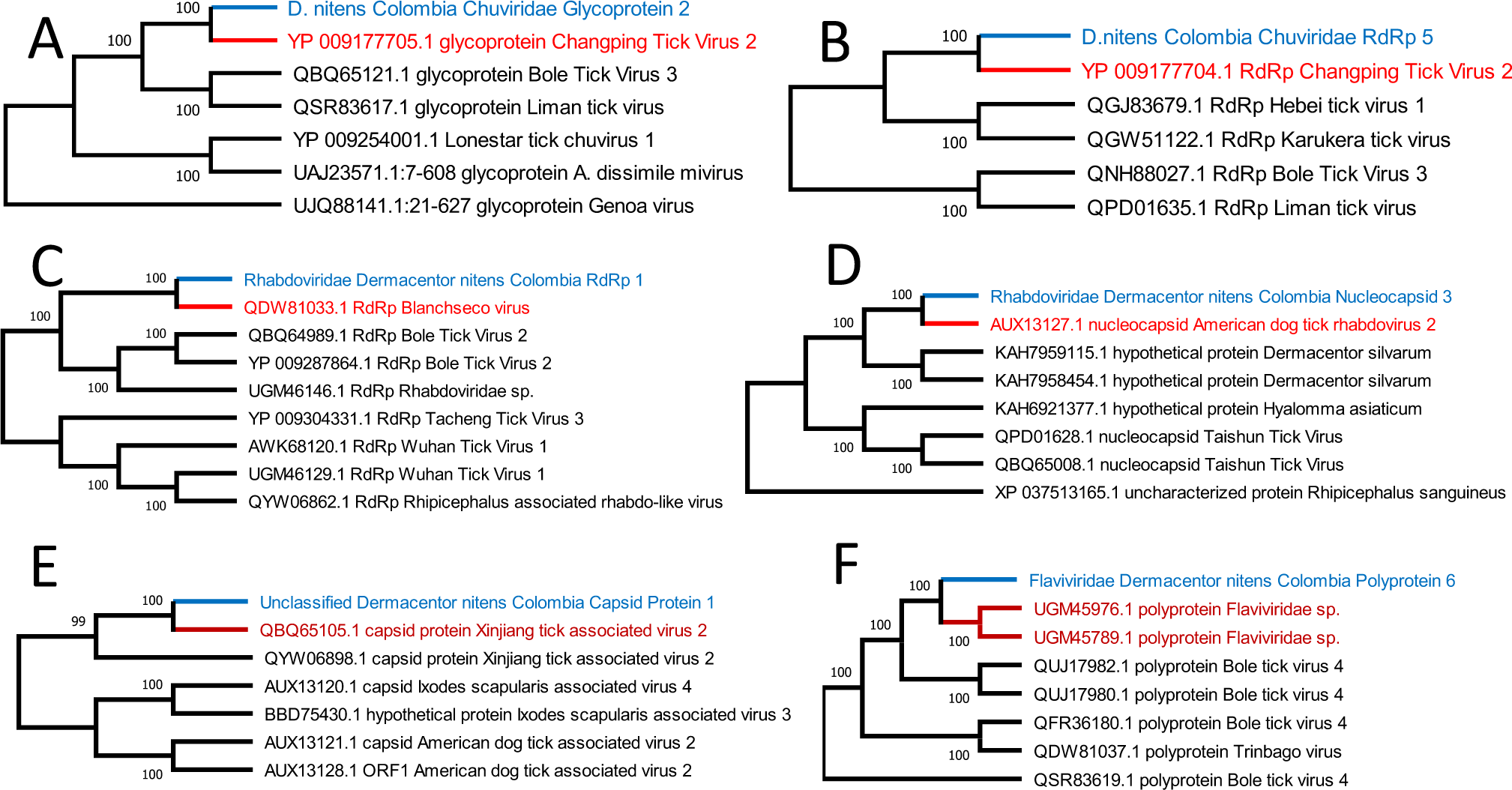
Phylogenetic relationship of the contigs for to RNA viruses captured in the *D. nitens* samples in this study. The maximum likelihood cladograms were constructed with complete deletion of assembly gaps. Bootstrapping percentages in 500 replications are shown at the nodes. The contig *D. nitens Colombia Chuviridae Glycoprotein 2* encodes a Glycoprotein gene with a length of 668 bp **(A)**, *D.nitens_Colombia_Chuviridae_Polymerase_5* encodes an RNA-dependent RNA polymerase with a length of 2156 **(B)**, *Rhabdoviridae_Dermacentor_nitens_Colombia_Polymerase_1* encodes an RNA-dependent RNA polymerase with a length of 7061 bp **(C)**, *Rhabdoviridae_Dermacentor_nitens_Colombia_Nucleocapsid_3* encodes a nucleocapsid with a length of 524 bp **(D)**, *Unclassified_Dermacentor_nitens_Capsid_Protein_1* encodes a capsid protein with a length of 168 bp **(E)**, *Flaviviridae_Dermacentor_nitens_Colombia_Polyprotein_6* encodes a polyprotein with a length of 5140 bp **(F)**. Names in blue correspond to the viral contigs found in this study, and red names correspond to the closest viral protein sequence in the GenBank database. The GenBank accession numbers are shown at the beginning of the names of taxa.

## 4 Discussion

Hard ticks harbor a considerable diversity of bacteria and viruses, of which there are significant pathogens to humans or domestic animals (2,4,6,8,61–63). A comprehensive survey of tick microorganisms may allow us to uncover the spectrum of the vectorial capacity of ticks for known pathogens and yield novel potential pathogenic microorganisms. In addition, it may provide a better understanding of the interactions among microorganisms under different environmental conditions. Thus, identifying symbiotic microorganisms and their effects on the vectorial capacity is critical for predicting future outbreaks caused of febrile diseases of unknown etiology (3). In this study, metatranscriptome and bacterial 16S rRNA sequencing enriched the sequence database with newly uncovered *Francisella*-like Endosymbionts (FLE) and virus genes in the blood-fed *D. nitens* originating from three different geographical areas in Colombia.

Differences in the bacterial compositions of ticks collected from animals coming from Bolivar, Antioquia, and Cordoba populations were found in either inclusion or exclusion of the FLE sequences. (Figures 1C and 1D). The NMDS plot for 16S sequences revealed clusters for tick geographical origin with a unique bacterial assortment. Geographically separated populations of ticks have previously been shown to have distinctive microbial compositions in a number of tick species (17,39,40,64,65). Microbial compositions could be influenced by other factors, such as the degree of tick engorgement, which has been reported previously (66–68). The capacity of ticks to acquire and spread pathogens may be significantly impacted by these variations in the microbial composition.

We found that the most abundant bacterium was FLE (80% of classified reads), which is phylogenetically related to the pathogenic bacteria *F. tularensis*, and causes tularemia in humans (9). While *Dermacentor variabilis* and *Dermacentor andersoni*, are known to carry this pathogen and are common in the northern hemisphere, the effect of FLE interaction with pathogens and their role in disease transmission remain unknown (1,11,17,69,70). Previous results have shown a positive association of vertically-transmitted FLE against pathogenic *Francisella novicida* artificial infection in *D. andersoni*, however *F.novicida* is not considered a tick-borne pathogen, which means this interaction is unlikely to happen in natural conditions (7).

Our result shows that the microbial composition of *D. nitens* appears to vary depending on the geographic location of the species’ population. We observed overall higher proportion of FLE compared to those previously reported in *D. variabilis* (62%), and *D. occidentalis* (41%) in the Americas (17,71). This highly abundant FLE was in accordance with previous 16S rRNA sequencing studies on whole-body samples obtained from partially or fully-engorged adult *Dermacentor* spp., females as *D. variabilis*, *D. marginatus*, *D. reticulatus*, *D. silvarum*, and *D. albipictus* (71–74). Metatranscriptomic analysis suggested high levels of FLE coverage (*i.e.*, transcript per million reads TPM) for Cordoba samples, but without statistical significance in all pairwise comparisons by Student t-test. 16S rRNA analysis, showing the relative abundance, also suggested that the Cordoba population is richer in FLE. The department of Cordoba, an agricultural stronghold in northern Colombia, has a constant flow and exchange of animals. Thus, the associated ticks may be exposed to a more diverse bacterial environment, which may explain the increased detection frequency of main endosymbiont and transient bacteria, through mechanisms such as horizontal transfer (1,64). These tendencies of small differences in the communities of endosymbionts related to the geographical origin of the ticks have also been reported for *D. occidentalis* (17). In other tick species, such as *Ixodes scapularis*, the endosymbiont population has been shown to impact pathogen infection processes. An unaltered intestinal microbiota favored colonization of *Borrelia burgdorferi s.l.*, while an induced microbial dysbiosis environment showed a negative effect by blocking colonization of *Anaplasma phagocytophilum* (1,19). In *D. nitens*, the transmission of human pathogens is yet unknown; however, *D. nitens* ticks collected from equines in Brazil were found positive for *B. burgdorferi s.l.*, the complex known as the causal agent of Lyme disease in the Americas (75). While *D. nitens’* potential as a Lyme disease vector, and the roles of FLE population have not been documented, the initial characterization of FLE population, may provide insights into their involvement in tick vector competence.

Our FLE sequence analysis revealed three different *D. nitens* FLE variants, OTU001, 002, and 010, with relatively large variations (8 to 21 bp or 1.7 to 4.5% difference) in the V3-V4 region. The source of these variants are likely from different strains that occurs in all three geographical locations. While the genus *Francisella* contains three 16S rRNA copies, we exclude the possibility of intra-genomic variations from these copies based on a study that described 99.65% minimum similarity average in 1374 Proteobacteria genomic sequences of 16S rRNA (76). These results are comparable to our previously reported study in *Amblyomma americaun*, where at least two different strains of *Coxiella*-like endosymbionts were found, at the individual tick level (44). Three *D. nitens* FLE OTUs were monophyletic and clustered while this cluster is also grouped with the FLE of other *Dermacentor* FLEs (Figure 2). However, FLEs of *R. microplus* and *I. scapularis* were also grouped in this clade (77), indicating, first that endosymbionts are more diverse than previously thought, and second that relatively recent independent invasions or transfers of FLEs frequently occurred, as it has been shown that the FLE initially evolved from the pathogenic *Francisella* species (1,12,13,59,60,62,71,77,78).

Metatranscriptomics revealed several contigs highly similar to viral families. Rhabdoviridae family was found as the most abundant and common in the pools of all sequences. This group of Rhabdoviridae viruses (Figure 4D) were also reported for different Ixodidae species such as *Rhipicephalus annulatus*, *R. sanguineus*, *Hyalomma marginatum*, *H. asiaticum*, and *D. variabilis* in the United States (23,24,26). Blanchseco virus (Rhabdoviridae family) was found in one pool of *Amblyomma ovale* ticks infesting cattle and dogs in Trinidad and Tobago (27). Similarly, we have identified Chuviridae-related sequences in the *D. nitens* RNA pools as the second predominant viral family (Figure 4A). Chuviridae is a newly-proposed viral family, that constitutes a large monophyletic group, clustering in an intermediate phylogenetic branch between segmented and unsegmented negative-sense RNA viruses identified in ticks, true flies, mosquitoes, cockroaches, and crabs (23). The most closely related to the *D. nitens* virus found in this study was previously identified in China (Figure 4A) with 90.2% (11,275 out of 12,500 bp) nucleotide sequence identity. The similar viruses in different continents may originate from historical commerce of animals.

We found geographical differences in the Rhabdoviridae family according to the contig Rhabdoviridae_RdRp that showed differences between Antioquia and Cordoba regions (p = 0.02), and the sequence coverage for Rhabdoviridae_Nucleocapsid is predominant in Bolivar when compared with those in other two regions (p = 0.03). The frequency data support unique viral compositions in different region (Supplementary Table 4). The coverage of the viral gene composition among the ticks in three different populations showed statistical differences in transcripts classified into the Rhabdoviridae family (Supplementary Table 5). A previous study with *R. microplus*, *D. nitens*, and *R. sanguineus s.l.* in the Magdalena Valley and Magdalena/Urabá ecoregions in Colombia reported the presence of Flaviviridae, Rhabdoviridae, Chuviridae, and Unclassified viruses (29). We conclude that the core RNA virome composition appears to be poor compared with the bacterial endosymbiotic communities. However, identifying viruses by using preexisting viral sequences in the GenBank may be limited for the discovery of novel viruses. This sequence-based survey needs further investigation to understand whether those are transiently acquired with the mammalian blood or established and vertically transmitted.

Overall, this study offers a description of the diversity of bacterial and viral communities of partially-fed *D. nitens* female ticks collected in animals originating from three Colombian regions based on our 16S rRNA sequences and transcriptomic analysis. In addition to the differentiated geographical populations in the bacterial and viral composition, we also found multiple co-existing strains of FLE and six different viruses in *D. nitens,* which offers the foundation for future studies. A deeper understanding of the microbial and viral communities hosted by ticks can be utilized to develop future measures to mitigate tick pathogen transmission.

## 5 Conflict of Interest

The authors declare that the research was conducted in the absence of any commercial or financial relationships that could be construed as a potential conflict of interest.

## 6 Author Contributions

Conceptualization, BL-R, and YP; experimental design—AH-R, LPM-R, BL-R, and YP; sample collection— AH-R, GMV, HA, AT-C, and GV-T; sample processing— AH-R; data analysis—AH-R, AC-T, and YP; writing—original draft preparation, AH-R, and YP; writing—review and editing, AH-R, LPM-R, KS, GMV, MLF, YP, and BL-R; funding acquisition, GMV, MLF, YP, and BL-R. All authors have read and agreed to the published version of the manuscript.

## 7 Funding

This study was partially funded by the Armed Forces Health Surveillance Division (AFHSD), Global Emerging Infections Surveillance (GEIS) Branch, PROMIS ID 2019 P0143_19_N6_04, 2020 P0144_20_N6_04 to MLF and GMV, USDA-multistate fund KS17MS1443-NE1443 to BL-R, NIH-NIAID R21 AI163423 and USDA-NIFA GRANT13066347 to YP.

## 8 Disclaimer

The views expressed in this article reflect the results of research conducted by the authors and do not necessarily reflect the official policy or position of the Department of the Navy, Department of Defense, nor the U.S. Government.

## 9 Copyright statement

Some authors of this manuscript are employees of the U.S. Government. This work was prepared as part of their official duties. Title 17 U.S.C. §105 provides that “Copyright protection under this title is not available for any work of the United States Government”. Title 17 U.S.C. §101 defines a U.S. Government work as a work prepared by a military service member or employee of the U.S. Government as part of that person’s official duties.

## 10 Acknowledgments

The authors thank the employees of “La Rinconada” slaughterhouse, the collaborators from Universidad de Antioquia, the Kansas State Department of Entomology, and the College of Agriculture.

## Notes

### Competing Interest Statement

The authors have declared no competing interest.

